# Cell body clustering drives gap junction-mediated synchronous activity in command neurons

**DOI:** 10.64898/2026.02.26.708359

**Authors:** Kristen Lee, Josmarie Graciani, Natalie Rico Carvajal, Zhehao Zhu, Matt Q Clark, Chris Q Doe

## Abstract

The nervous system contains densely packed cell bodies, yet the role of neuronal cell body position in circuit function is poorly understood. Here we show that four *Drosophila* Moonwalker Descending Neurons (MDNs), command neurons for backward locomotion, must maintain cell body contact to allow gap junction-dependent synchronous activity necessary to initiate backward walking. MDNs express the transcription factor Hunchback, which drives expression of the Lar cell adhesion molecule; Hunchback, Lar, and its ligand Dlp promote MDN cell body clustering and backward walking. When clustered, the gap junction protein Inx8 allows synchronous firing of MDNs, which is required to initiate backward walking. These findings reveal a previously unappreciated role for cell body clustering and synchronous firing in neural circuit function.

## Main Text

Neuronal electrical activity must be precisely regulated to avoid excitotoxicity or failure to activate circuits (*1*). Proper neuronal communication allows the brain to integrate environmental cues and generate motor output. Neuronal circuits are often depicted as one neuron communicating with another, although neurons are rarely functioning alone. Typically, multiple neurons make up a neuronal type, which communicates with other neuronal types (i.e. other neuronal ensembles) (*2*). However, little is known about the principal features needed for neurons comprising a neuronal type to function properly within a neuronal network. Neuronal features such as morphology, neurotransmitter choice, and ion channel composition all have the potential to modify neuronal activity and connectivity (*3*). Here we add one more attribute to this list: cell body positioning to regulate electrical coupling.

To identify features regulating circuit activity, we used the well-characterized *Drosophila* Moonwalker Descending Neuron (MDN) escape circuit, which commands backward walking (*4*). There are four MDNs, two on each side of the midline, which respond to environmental stimuli before synapsing with neurons in the brain and ventral nerve cord (VNC) to initiate and maintain backward walking (*5–10*). The third thoracic segment of the ventral nerve cord primarily controls the hindlegs. Within this neuropil, MDNs major output is onto the pre-motor neurons LBL40 and LUI130 that drive backward walking (*8*).

The MDN circuit is uniquely equipped to investigate the relationship between neuronal activity and neuronal identity within a multi-neuron ensemble. We find a single transcription factor, Hunchback (Hb), is required in the four MDNs for expression of the Lar-Dlp cell adhesion complex, which generates a tight four-cell body cluster of MDNs. Strikingly, without cell body clustering, MDN-induced backward walking does not occur. This system allows us to address a novel question within the field of neuroscience: what role does cell body location have for neuronal circuit function? We find that cell body clustering of MDNs is necessary for synchronous firing of all four MDNs via gap junction electrical synapses, thereby identifying a previously undescribed feature of neuronal types necessary for network activity. It is possible that cell body location may be an important principal for neuronal network function across species; electrical synapses are well-documented to be present within populations of a single neuron type, including the fly lamina, worm AVA interneurons, and mammalian inferior olive neurons (*11–13*). Here, our data support a model where synchronous activity, enabled by clustered cell bodies, is required to translate diverse inputs into a temporally coherent burst of activity that is required for initiating backward walking.

### Hunchback is required in MDNs for backward walking

We previously showed that MDNs express Hb across larval life (*14*). Here, we show that MDNs maintain Hb expression into adulthood (Fig. 1A). As anticipated, expressing *UAS-Hb^RNAi^* produces a strong Hb knockdown in adult MDN neurons (Fig. 1A) (*15*). Hb knockdown in larval MDNs produced an increase in MDN-induced backward locomotion (Fig. 1B), due to an increase in synaptic connectivity with A18b, a downstream partner in the backward circuit (*14*). To determine the function of Hb in the adult MDN, we knocked down Hb in post-mitotic MDNs and assayed walking behavior. The adult MDN driver turns on during mid-metamorphosis.

**Figure 1.**
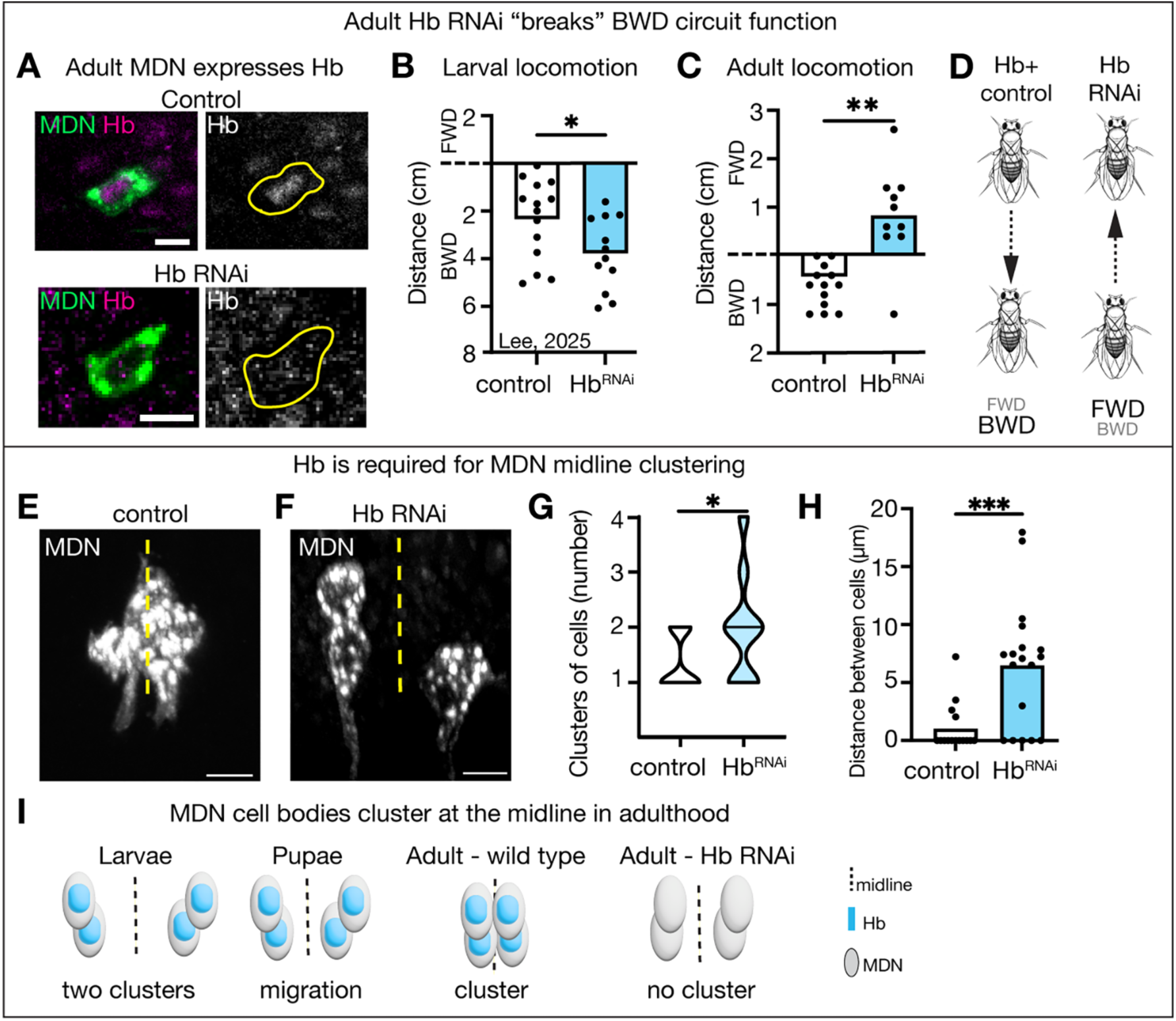
Hunchback is required for adult MDN behavior and cell body clustering at the midline. (**A**) Hb expression is lost when the Hb RNAi transgene is expressed in MDN. Scale bar, 5 µm. (**B**) Data from Lee, 2025(*14*); distance traveled when MDN is activated in larvae. (**C**) Distance traveled when MDN is optogenetically activated in adults. (**D**) Summary of behavior results. (**E-F**) Adult MDN cell body position relative to midline (dashed). Scale bar, 5 µm. (**G-H**) Quantification of E-F. (**I**) Cartoon representation of results. Statistical analyses were performed using unpaired two-sided t-test (*p<0.05, **p<0.01, ***p<0.001, ****p<0.0001).

In control animals, when adult MDNs are optogenetically activated, the animals walk backwards (Fig. 1C) (*4*, *16*). Unexpectedly, when Hb is knocked down in adult MDNs, optogenetic activation failed to elicit backward locomotion (Fig. 1C), thereby “breaking” the MDN circuit (Fig. 1D). Thus, Hb is regulating some adult MDN feature that is required to induce backward walking.

One possible explanation for the locomotor difference between larvae and adults would be neurotransmitter switching. The larval MDNs are excitatory cholingeric neurons (*4*, *17*), and Hb knockdown maintains ChAT expression and lacks GABA inhibitory neurotransmitter expression in adult MDNs (fig. S1A, B), indicating that Hb is regulating a different MDN feature required for backward walking. To determine if the MDNs have synapse abnormalities, we assayed MDN chemical synapse number. Loss of Hb did not alter MDN synapse number (fig. S1C-E). To determine if the MDNs have defective morphology, we assayed the morphology of single-labeled MDN neurons via Multi-Colored Flip Out (*18*). Loss of Hb did not alter MDN axon or dendrite morphology (fig. S2).

In larvae, two pair of MDNs are located off the midline; during metamorphosis the cell bodies move towards the midline before forming a tight four-cell cluster in adults (Fig. 1I; fig. S1C). Interestingly, Hb knockdown impaired clustering of MDN cell bodies at the midline (Fig. 1E-H). In contrast, Hb knockdown had no effect on MDN cell body position in the larvae (fig. S1F-H). We conclude that aggregation of MDN cell bodies at the midline during adulthood is disrupted in Hb knockdown (Fig. 1I).

### Hunchback promotes Lar expression, which is required for MDN midline clustering and backward locomotion

To determine whether loss of Hb or lack of MDN cell body clustering resulted in loss of MDN-induced backward walking, we screened for guidance cues that prevented midline clustering without altering Hb levels. We found that the Type IIa receptor-like protein tyrosine phosphatase Lar was expressed in MDNs during pupation and adulthood, but not in larvae (Fig. 2A-C). Next, we asked whether Lar was important for MDN cell body clustering. Expression of *UAS-Lar^RNAi^* in the adult MDN neurons resulted in a strong Lar knockdown (Fig. 2E, F) and disrupted midline clustering of MDNs (Fig. 2G-J) without altering Hb levels (t-test, p=0.842). We then asked if Hb was required for the expression of Lar; adult Hb knockdown in MDN resulted in significantly reduced Lar (Fig. 2D, F). We then assayed Lar knockdown to see if disruption of the MDN midline clustering, but not loss of Hb levels, resulted in failure to walk backwards upon optogenetic activation. Control animals walked backward when MDN was optogenetically activated, whereas Lar knockdown animals failed to go backwards (Fig. 2K). We conclude that Lar knockdown in MDNs “breaks” the backward locomotion circuit, without altering Hb levels.

**Figure 2.**
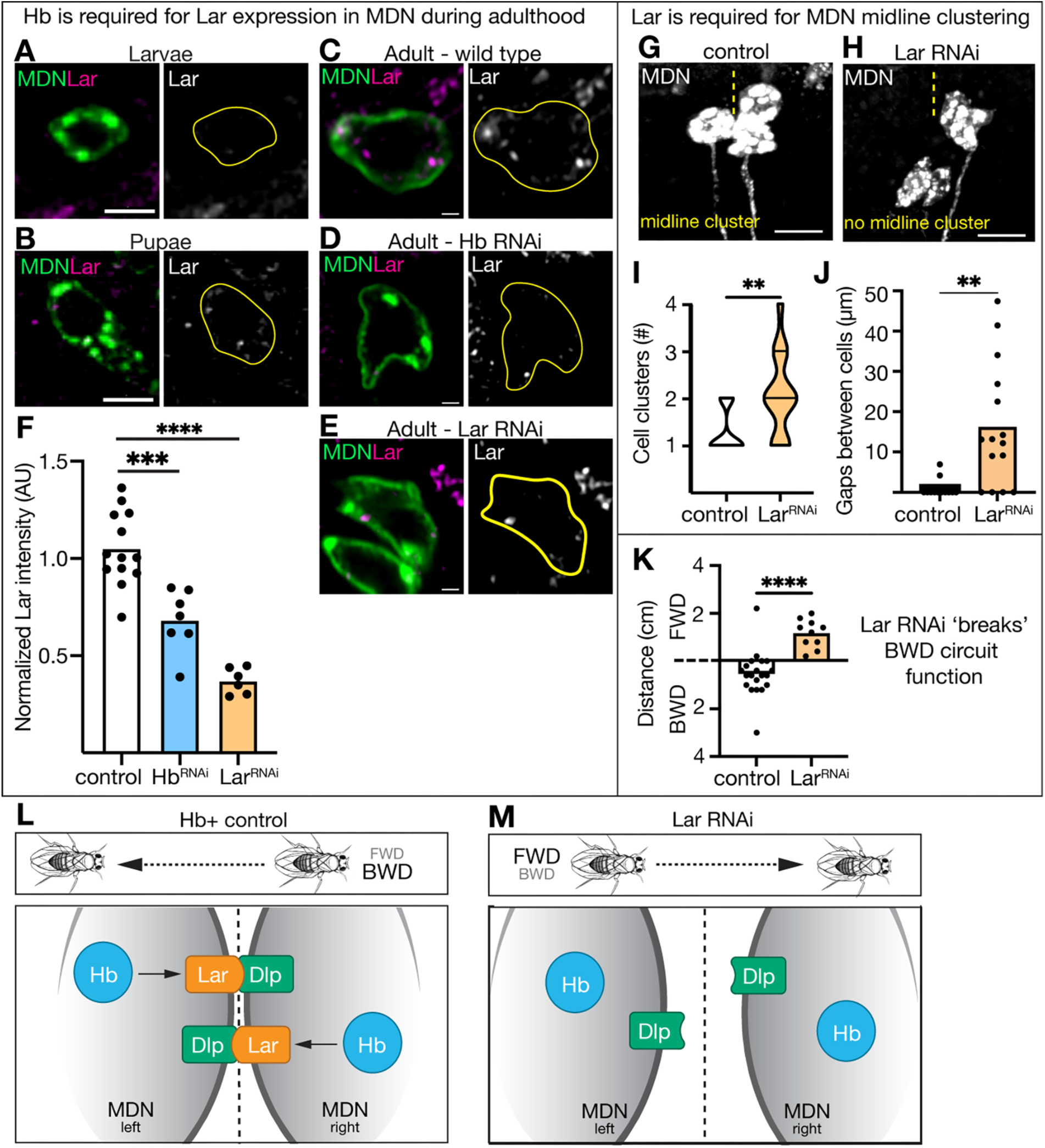
Hunchback promotes Lar expression, which is required for MDN midline clustering and backward locomotion. (A-C) Lar expression in MDN at indicated stages. A and B: scale bar, 5 µm. C: scale bar, 1 µm. (**D**) Lar expression is reduced when Hb RNAi is expressed in MDN. Scale bar, 1 µm. (**E**) Lar expression is reduced when Lar RNAi is expressed in MDN. Scale bar, 1 µm. (**F**) Quantification of C-E. (**G-H**) Adult MDN cell body morphology. Scale bar, 7 µm. Yellow dash line represents the midline. (**I, J**) Quantification of G-H. (**K**) distance traveled when MDN is optogenetically activated. (**L, M**) Cartoon representation of results. Statistical analyses were performed using a two-way ANOVA with Bonferroni’s multiple comparisons or an unpaired two-sided t-test, as appropriate (*p<0.05, **p<0.01, ***p<0.001, ****p<0.0001).

Lar has well-characterized roles in neuronal circuit wiring (*19*). Our data suggest a novel role of Lar: attracting neuronal cell bodies to the midline. To further demonstrate that Lar is required for MDN cell bodies to cluster at the midline, we assayed Lar ligands: N-cadherin (CadN), Sticks and stones (Sns), Syndecan (Sdc), and Dally-like protein (Dlp) (*19–22*). Like Lar, Dlp was expressed in pupal and adult MDNs but not in larvae (fig. S3A-C). Additionally, we investigated whether Dlp was important for MDN cell body clustering. We observed a moderate Dlp knockdown when expressing *UAS-Dlp^RNAi^* in the adult MDN neuron (fig. S3E, F), yet the partial Dlp knockdown still prevented MDN midline clustering without altering Hb levels (fig. S2G-J; t-test, p=0.691). Conversely, knocking down Hb in adult MDNs did not alter Dlp expression (fig. S3D, F). Taken together, these data lead to a linear model: Hunchback promotes Lar expression, Lar promotes MDN midline clustering by binding to Dlp, and MDN clustering is required for MDN-induced backward locomotion (Fig. 2L, M).

### Innexin 8 is required for MDN clustering and initiation of backward walking

The requirement of MDN cell body clustering for backward walking was surprising. How might disrupting the MDN soma cluster lead to failed backward walking induction? One possibility is that all four MDNs are gap junction coupled, resulting in adhesion between the cell bodies and synchronous firing. It is well-documented that electrical synapses can occur between cell bodies, regulate adhesion, and allow neurons to fire synchronously (*23*, *24*).

We assayed the widely expressed gap junction component Innexin 8 (Inx8; Flybase: ShakB) (*25*). We confirmed that Inx8 is detected in adult MDNs, specifically between MDN cell bodies and between primary neurites (Fig. 3A), with little at axon terminals (fig. S4A). We then tested whether Hb was required for Inx8 expression. We found that Hb knockdown did not remove Inx8 expression in MDNs but led to an increased number of smaller Inx8 puncta compared to control (Fig. 3B-F; see Discussion). We observed a similar result when Lar was knocked down (fig. S4B-E). Next, we assayed the role of Inx8 in MDNs. We expressed *UAS-Inx8^RNAi^*specifically in the MDNs, as well as assaying a homozygous viable *Inx8* null mutant. In both experiments we found significantly reduced Inx8 expression on MDNs (Fig. 3D-F), suggesting that MDN-MDN gap junctions are reduced or eliminated in these genotypes, which inhibited MDN cell body clustering (Fig. 3G-L). Importantly, MDN-specific expression of a wild-type Inx8 transgene in the *Inx8* mutant background fully rescued MDN midline clustering (Fig. 3J-L). Lastly, we investigated whether Inx8 knockdown disrupts backward walking upon MDN optogenetic activation. In this experiment, control animals went backward when MDNs were activated, whereas Inx8 knockdown animals paused (Fig. 3M). Given that Inx8, Lar, and Dlp are all cell surface molecules, we investigated whether they are associated on MDNs cell membrane. We found that Inx8 puncta were associated with both Lar and Dlp (Fig. 3N). We conclude that Inx8, Lar, and Dlp are interacting on MDN cell bodies to promote MDN clustering, which is required for proper behavioral output (Fig. 3O, P). We conclude that Innexin 8 is required for MDN clustering and initiation of backward walking.

**Figure 3.**
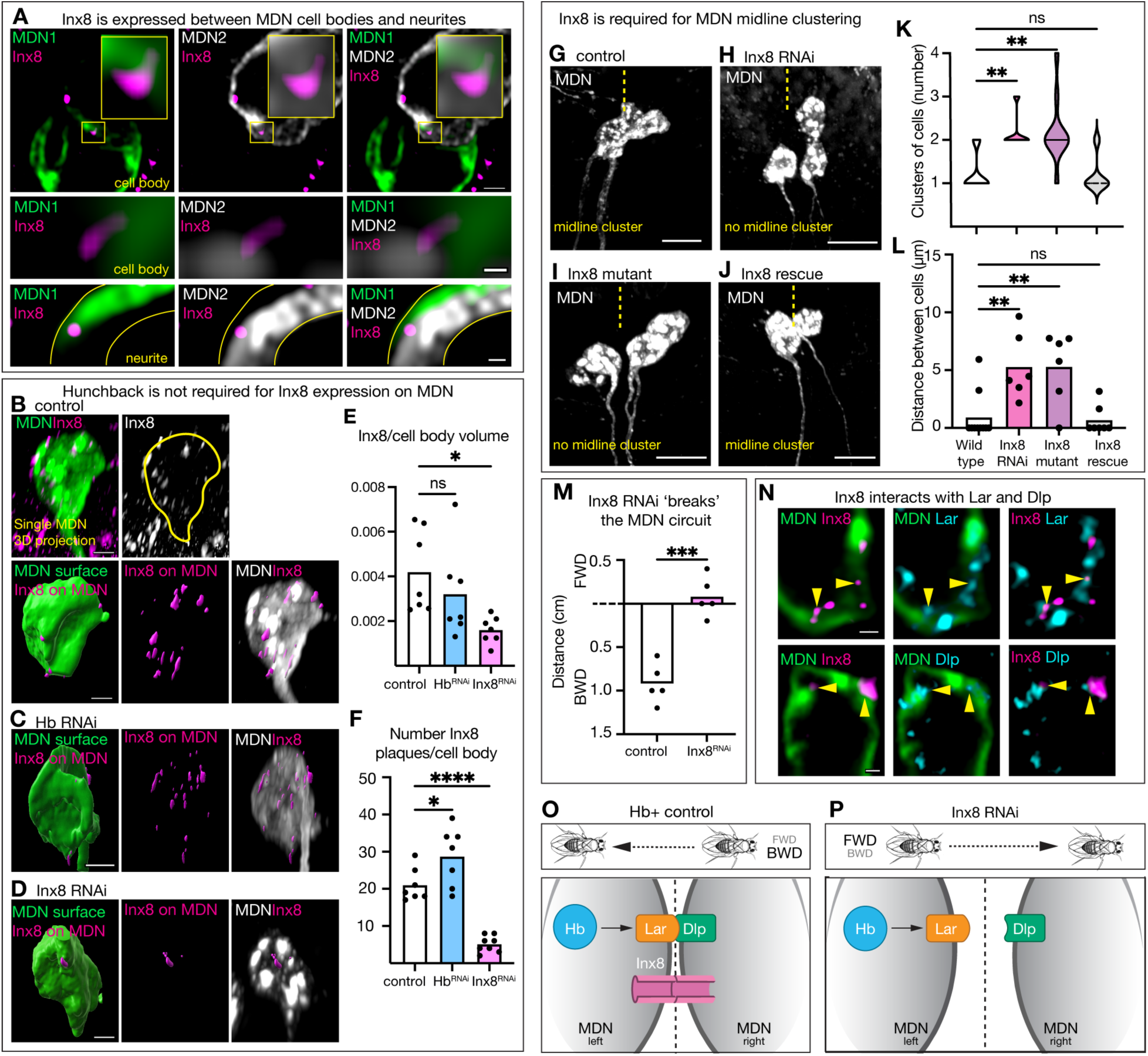
Innexin 8 is required for MDN midline clustering. (**A**) Inx8 is present between individual MDN cell bodies and neurites. Top: scale bar, 1 µm. Middle: scale bar, 0.1 µm. Bottom: scale bar, 0.3 µm. (**B–D**) Inx8 puncta expression on MDN cell bodies when Hb RNAi and Inx8 RNAi is expressed in MDN. Scale bar, 3 µm. (**E, F**) Quantification of B-D. (**G-J**) Adult MDN cell body morphology. Scale bar, 10 µm. Yellow dash line, midline. (**K, L**) Quantification of G-J. (**M**) Distance traveled when MDN is optogenetically activated. (**N**) Inx8 expression alongside Lar (top) and Dlp (bottom) on MDN cell bodies. Scale bar, 0.5 µm. (**O, P**) Cartoon representation of results. Statistical analyses were performed using a two-way ANOVA with Bonferroni’s multiple comparisons or an unpaired two-sided t-test, as appropriate (*p<0.05, **p<0.01, ***p<0.001, ****p<0.0001).

### MDN clustering is required for synchronous firing

We hypothesized that synchronous activity of the four MDNs is required for MDN-induced backward locomotion. We tested for synchronous firing in control, Hb knockdown, and Inx8 knockdown animals using the genetically encoded voltage indicator ASAP5 (*26*). We expressed the red-light gated cation channel, CsChrimson, in MDNs, and used two-photon microscopy to activate a single MDN. We then measured the voltage response of both the stimulated MDN and an adjacent non-stimulated MDN (typically 2 total cells; Fig. 4A). For each animal we calculated the likelihood of the non-stimulated MDN to fire synchronously with the stimulated MDN; we found that control MDNs were significantly more likely to have a synchronous voltage response compared to Hb and Inx8 knockdown animals (Fig. 4B). Overall, control non-stimulated cells frequently fired synchronously with the stimulated cell (Fig. 4C), while the non-stimulated Hb and Inx8 knockdown MDNs had much smaller voltage fluctuations and infrequent synchronous activity (Fig. 4D-E; population quantification in fig. S4F-H).

**Figure 4.**
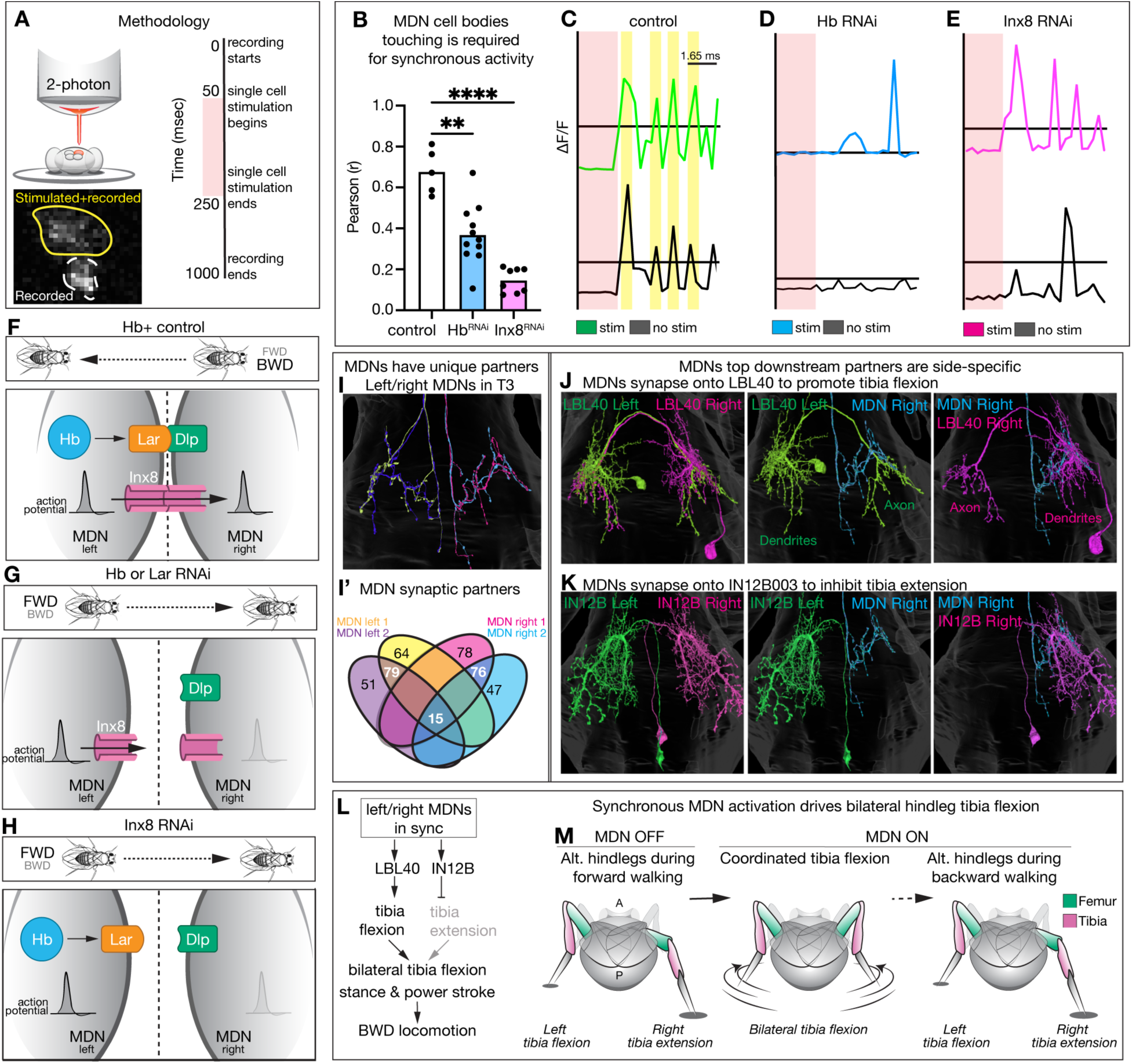
MDN clustering is required for synchronous firing, which drives initiation of backward walking. (**A**) Methodology for the genetically encoded voltage indicator experiments. (**B**) Pearson correlation coefficients of the likelihood adjacent MDN will fire when the stimulated MDN fires. Statistical analyses were performed using a two-way ANOVA with Bonferroni’s multiple comparisons (*p<0.05, **p<0.01, ***p<0.001, ****p<0.0001). (**C-E**) Voltage indicator traces between stimulated cell (colored) and adjacent cell (black) in a single animal when Hb or Inx8 RNAi is expressed in MDN. Yellow bands demarcate synchronous activity. Black horizontal lines represent resting membrane potential. (**F-H**) Cartoon representation of results. (**I**) Each individual MDN, and its downstream partners, is represented by a unique color. (**J**) Right MDN only forms synapses with the right LBL40, not the left. (**K**) Right MDN only forms synapses with the right IN12B003, not the left. (**L**) Neuronal circuit downstream of all four MDNs firing synchronously. (**M**) initiation of backward walking requires coordinated bilateral hindleg flexion.

Additionally, Hb knockdown in MDNs had a lower resting membrane potential, likely due to increased electrical coupling with neighboring inhibitory GABAergic cells (*27*) (fig. S4I-K). Taken together, we conclude that MDN clustering is required for synchronous firing and regulating membrane potential, which is required for MDN-induced backward walking (Fig. 4F-H).

### Synchronous MDN activity is required for initiation of backward walking

How does failure of MDN to fire synchronously result in a loss of backward locomotion? In wild type, initiation of backward locomotion is triggered by a brief, bilateral hind leg flexion, followed by a bilateral backward power stroke, before alternating leg movements characteristic of backward walking (*8*, *10*). We hypothesized that loss of synchronous MDN activation might generate an uncoordinated hindleg flexion, preventing backward walking initiation. This model predicts that the four MDNs have a diverse population of output neurons that require synchronous MDN firing to generate robust, bilateral hind leg movement. Consistent with this hypothesis, we identified MDN downstream partners using the Male Adult Nerve Cord connectome (*28*), and found that the four MDNs had very few common outputs (3.6% or 15 total), with hundreds of outputs targeted by individual MDNs or left/right pairs (Fig. 4I, I’). The neurons with the most synapses were the premotor neurons LBL40 and IN12B003 (Fig. 4J, K), which are unique to MDN left/right pairs (Fig. 4I’). LBL40 activation triggers hindleg flexion, whereas IN12B003 inhibits hindleg extension (*8*, *10*). Even these neurons have diverse outputs: LBL40 targets the contralateral neuropil, while IN12B003 remains on the ipsilateral side, highlighting how synchronous activation of the left/right MDNs is needed to activate bilateral tibia flexion while suppressing bilateral tibia extension (Fig. 4L, M). In our experiments, knockdown of Hb, Lar, or Inx8 abolishes MDN synchronous firing and fails to initiate backward walking. We propose that loss of MDN synchronous activation results in the uncoordinated activity of the diverse population of MDN output neurons, preventing coordinated bilateral tibia flexion and failure to initiate backward locomotion.

## Discussion

Our findings highlight the importance of cell body position for neuronal circuit function. We find that anatomical location of cell bodies and expression of ligand-receptor cell adhesion proteins, along with gap junction electrical synapses, are required for MDNs to initiate backward walking. Notably, these functions are independent of MDN chemical synapses and axon/dendrite morphology. The transcription factor Hb activates expression of the ligand-receptor pair, Lar-Dlp, which is required for the MDNs to form a single cell body cluster at the midline (Fig. 1, 2). The four MDN cell bodies form electrical synapses with each other, allowing them to fire synchronously (Fig. 3, 4). Disrupting cell body location by knocking down Hb, Lar, or Inx8 inhibits MDN-induced backward locomotion (Fig. 1-3). Moreover, we show that each MDN forms downstream connections with a mostly non-overlapping pool of neurons. Thus, MDN synchronous activation is required to coordinately activate each of the “micro-circuits” downstream of MDN, which are essential for backward walking (Fig. 4)(*8*).

Our observation that Hb promotes neuronal cell body adhesion to allow synchronous firing is a novel result. Hb is a well-characterized transcription factor that generates early-born neuronal identity in *Drosophila* (*29*) and mammalian cortex and retina (*30*, *31*). In the *Drosophila* larvae, Hb is required for post-mitotic MDN morphology and chemical synapse number (*14*). Here we show that during adulthood, Hb plays no role in MDN axon/dendrite morphology (fig. S2) or chemical synapse number (fig. S1C-E).

We find that Hb actives expression of the cell adhesion molecule Lar in pupal and adult MDNs (Fig. 2). During pupal stages, the four MDN cell bodies require Lar-Dlp to cluster together. Outside of the CNS, Lar plays a role in cell migration in the follicular epithelium (*32*, *33*). Additionally, we may have uncovered a new function of Lar in supporting electrical synapse plaque size. Proteomic work in Zebrafish shows that cell adhesion molecules are in close proximity to Mauthner electrical synapses (*34*), but the relationship between electrical synapses and cell adhesion molecules has not been tested *in vivo*. Here we find that MDN electrical synapses are in close proximity with the cell adhesion molecules Lar and Dlp, and we show that reducing Lar expression (via Hb or Lar knockdown) leads to smaller electrical synapse puncta on MDN (Fig. 3; fig. S4), a novel result within the gap junction field. These data support the hypothesis that the transmembrane protein Lar is required for Inx8 organization at MDN-MDN contact sites. The likelihood that cell adhesion molecules are required for gap junction organization or stability is provocative; to date, most *in vivo* experiments have only characterized adherens junctions and scaffold protein interactions with gap junctions (*24*, *35*). Given that fly innexin gap junctions can function in vertebrate cells, and vertebrate connexin gap junctions can function in invertebrates (*36*, *37*), these results are likely translatable to other systems.

Although gap junctions are found in non-neuronal cells (*38*, *39*), electrical synapses are present in escape circuits across species. In *C. elegans*, electrical synapses are present between the left and right AVA interneurons, which command forward and backward locomotion (*40*). In this system, electrical coupling is required for asymmetric inhibitory and excitatory sensory input to be processed and integrated into the correct motor response (*13*, *40*) – a striking example of the importance of synchronous activity between a neuron ensemble. The *Drosophila* Giant Fiber neurons control a jumping escape response and forms electrical synapses with downstream motor neurons that are required for escape behavior (*41*, *42*). Interestingly, the transmembrane protein Frazzled is necessary for both Giant Fiber morphogenesis and gap junction development (*43*). The Mauthner cell in Zebrafish initiates a fast C-bend escape behavior and has pre- and post-electrical synapses with upstream interneurons (*44*).

The left/right Mauthner cells are strategically <u>not</u> synchronous in this neural network; unilateral firing allows precise directional turning away from aversive stimuli (*45*). Like the Giant Fiber and Mauthner cell circuits, most previous studies assaying the influence of gap junctions on behavior focus on electrical coupling between distinct cell types (*46–49*). These examples highlight the importance of neuronal ensembles being electrically coupled. Gap junctions are not formed promiscuously between random adjacent cells (*50*, *51*) and cAMP has been shown to be a crucial regulator of electrical synapse specificity (*52*). Our data not only add to this growing body of literature but demonstrate the absolute requirement of proper electrical coupling within a neural network for behavior: without synchronous activity of the four MDNs, the fly is incapable of initiating backward walking.

Previous work used a genetic technique to activate left or right MDNs, finding that asymmetric activation still initiated backward walking but skewed the fly to turn to the contralateral side (*5*). We believe that their results complement our study; here, we find that each individual MDN forms synapses with diverse populations of downstream neurons. Therefore, due to the electrical coupling between MDNs, there is a mechanism defining how asymmetric activation of MDN could elicit backward walking. Sen et al (*5*) theorized “each of the four MDNs can act independently and that collectively they control both the magnitude and direction of backward locomotion”.

Our data demonstrate that MDNs do not work independently. The majority of the shared neurons downstream of MDN are side-specific (Fig. 4), and Cheong et al (*10*) proposed that side-specific inhibition of a front leg interneuron downstream of MDN likely contributes to turning. We agree that all four MDNs collectively control the initiation of backward walking. Here, even though all four MDNs are exposed to light for optogenetic activation at the same time, cell body clustering is required for normal circuit output. We believe a combination of factors contribute to this - such as resting membrane potential differences, gap junction regulation of response magnitude, and coordinated downstream circuit activation – demonstrating the requirement of coupling between the four MDNs (fig. S5). Taken together, our study has implications not just in the understanding of *Drosophila* motor output, but also in recognizing cell body location as a novel regulator of neural output.

## Acknowledgments

Antibodies obtained from the Developmental Studies Hybridoma Bank, created by the NICHD of the NIH and maintained at the University of Iowa, Department of Biology, Iowa City, IA, were used in this study. Stocks obtained from the Bloomington Drosophila Stock Center (NIH P40OD018537) and Vienna Drosophila Resource Center were used in this study. We thank the Desplan lab (NYU) and the Clandinin lab (Stanford) for reagents. We also thank Adam Fries (UO Microscopy Core) for help designing the 2-photon experiments, Adam Miller and lab members (UO) for project feedback, and Heather Pollington (UO) for help making all the schematics used in the figures. This article is subject to HHMI’s Open Access to Publications policy. HHMI lab heads have previously granted a nonexclusive CC BY 4.0 license to the public and a sublicensable license to HHMI in their research articles. Pursuant to those licenses, the author-accepted manuscript of this article can be made freely available under a CC BY 4.0 license immediately upon publication.

## Author Contributions

Conceptualization: KML, CQD Data curation: KML, JG, NRC, ZZ

Data analysis: KML, JG, NRC, ZZ, MC Visualization: KML, CQD

Funding acquisition: KML, CQD Supervision: KML, MC, CQD Writing – original draft: KML

Writing – review & editing: MC, CQD

## Data, code, and materials availability

All data will be deposited in databases for open access, and/or by request.

## Supplementary Materials for

### Materials and Methods

#### Fly husbandry

All flies were reared in a 25C room at 50% relative humidity with a 12 hour light/dark cycle. All flies used in experiments were 4-5 day old adults reared on standard cornmeal food medium. All comparisons between groups were based on studies with flies grown, handled, and tested together.

#### Fly strains

Short name, source, and full genotype of each individual fly strain used in this study is outlined below. Detailed genotypes for each experiment are provided in the corresponding methods section.

MDN-Gal4 (Bidaye, 2014^1^): VT50600.p65AD (attp40)/CyO; VT44845.Gal4DBD(attp2)/TM3,Ser or SS03131-Gal4

Control RNAi (BDSC# 31603):;; UAS-Luc RNAi TRiP.JF01355 (attp2) Hb RNAi (BDSC# 34704):;; UAS-Hb RNAi TRiP.HMS01183 (attp2) Lar RNAi (BDSC# 40938):; UAS-Lar RNAi TRiP.HMS02186 (attp40); Dlp RNAi (BDSC# 50540):; UAS-Dlp RNAi TRiP.GLC01658 (attp40); Inx8 RNAi (BDSC# 27292):;; UAS-shakB RNAi TRiP.JF02604 (attp2)

Inx8 mutant (Desplan Lab): ShakB2;;

Inx8 rescue (Desplan Lab): ShakB2; UAS-ShakB; UAS-ShakB Chrimson (BDSC# 55134): UAS-CsChrimson.mVenus (attp18);;

MCFO (BDSC# 55134 and 77140): hs-FlpG5::Pest; 10X UAS(frt.Stop)myr::smGDP-V5-THS-10XUAS(frt.Stop)myr::GDP-Flag;

ASAP5 (Clandinin Lab and BDSC# 55134): UAS-Chrimson::mVenus (attp8); UAS-ASAP5/CyO; TM2/TM6

LBL40-LexA (Feng, 2020^2^):; VT021418-LexAGADPI (attp40);

*Reporters used* (BDSC# 64092): LexAop-myr::smGdp-V5, UAS-myr::smGdp-HA;;

*Reporters used* (BDSC# 32197):;;UAS-myr::GFP

#### Hb, Lar, Dlp, and Inx8 knockdown experiments

The following genotypes were utilized for these experiments.

Control knockdown genotype: LexAop-myr::smGdp-V5, UAS-myr::smGdp-HA; VT50600.p65AD; VT44845.Gal4DBD/ UAS-Luc RNAi TRiP.JF01355.

Hunchback knockdown genotype: LexAop-myr::smGdp-V5, UAS-myr::smGdp-HA; VT50600.p65AD; VT44845.Gal4DBD/ UAS-Hb RNAi TRiP.HMS01183

Lar knockdown genotype: LexAop-myr::smGdp-V5, UAS-myr::smGdp-HA; VT50600.p65AD/UAS-Lar RNAi TRiP.HMS02186; VT44845.Gal4DBD

Dlp knockdown genotype: LexAop-myr::smGdp-V5, UAS-myr::smGdp-HA; VT50600.p65AD/UAS-Dlp RNAi TRiP.GLC01658; VT44845.Gal4DBD

Inx8 knockdown genotype: LexAop-myr::smGdp-V5, UAS-myr::smGdp-HA; VT50600.p65AD; VT44845.Gal4DBD/ UAS-shakB RNAi TRiP.JF02604

#### Innexin 8 mutant and rescue experiments

The following genotypes were utilized for these experiments. Innexin 8 mutant: ShakB2; VT50600.p65AD; VT44845.Gal4DBD/UAS-myr::GFP Innexin 8 rescue: ShakB2; VT50600.p65AD/UAS-ShakB; VT44845.Gal4DBD/UAS-myr::GFP

#### Immunohistochemistry

Standard confocal microscopy and immunocytochemistry methods were performed. In short, brains from 4-5 day old female flies were dissected in ice cold hemolymph-like buffer and fixed with 4% paraformaldehyde for 40 minutes, depending on age. The tissue was exposed to normalized donkey serum block for either 40 minutes at room temperature or overnight at 4C. After, the tissue was exposed to a primary antibody solution overnight at 4C. After being washed with 0.3% PBST, samples were exposed to a secondary antibody solution overnight at 4C. After a series of dehydration, tissue was mounted with DPX.

Primary antibodies used: Rabbit anti-Hunchback (1:400; Doe lab); Rat anti-HA (1:100; Sigma #11867423001; RRID AB_2687407); chicken anti-V5 (1:800; Fortis Life Sciences A190-118A; RRID AB_66741); mouse anti-Flag (1:1000; Sigma F1804; RRID AB_262044); Mouse anti-Lar (1:50; Developmental Studies Hybridoma Bank 9D82B3; RRID AB_528202); Mouse anti-Dlp (1:100; Developmental Studies Hybridoma Bank 13G8; RRID AB_528191); Guinea pig anti-Inx8 (1:1000; Desplan lab); rabbit anti-GABA (1:200; Sigma A2052; RRID AB_477652). Secondary antibodies were from Jackson ImmunoResearch (Donkey, 1:400).

#### Image acquisition and processing

Confocal image stacks were acquired on a Zeiss LSM900 Airyscan 2 microscope using a Plan-Apochromat 63x/1.4 Oil DIC M27 objective. Nyquist sampling, a pinhole of 1 Airy Unit, and optimal Z-stack step sizes were used. The imaging parameters were optimized to fill the detector’s dynamic range while avoiding pixel saturation.

Super resolution Airyscan imaging was done for all Lar, Dlp, and Inx8 staining. For these images, the scan zoom was set to 1.3X and pixel scaling was set to 0.043 µm X 0.043 µm X 0.170 µm. All images were obtained with a frame averaging of 4 and a pixel dwell time of approximately 1.15 µsec. All images were processed either in Fiji (https://imagej.new/fiji) or Imaris version 10.2 (https://imaris.oxinst). Methodology for specific analyses using these software packages are described below. Figures were made using Adobe Illustrator.

#### Quantification of cell body clusters

A cell body cluster was defined as 2 or more cell bodies that were touching. The number of cell body clusters were manually counted.

The distance between cell body clusters was calculated by averaging the shortest distances between all the cell body clusters in a single animal.

#### Quantification of pixel intensity

All pixel intensity quantification was done manually using the “Measure” feature in Fiji. The freehand tool was used to outline the cell body. Measurements were set to “Area” and “Raw Integrated Density”. A middle slice of the total cell body was measured. For each cell body, the “Raw Integrated Density” was divided by the “Area”. The sum of these values is reported.

#### Quantification of Innexin 8 puncta

Imaris image analysis software was used to put a surface over the MDN cell body and the Inx8 staining. The “surface on surface” feature was used to isolate all the Inx8 surfaces on the MDN cell body surface. By selecting the Inx8 on MDN surfaces, Imaris reported the total volume and total number of Inx8 puncta.

#### Larval optogenetic behavior

Larval behavior methodology was described previously^3^.

#### Adult optogenetic behavior

For adult optogenetic behavior experiments, all-trans retinal (ATR; Sigma-Aldrich Cat.# R2500) stock solution (100 mM) was prepared by dissolving 100 mg in 3.52 mL of 100% ethanol. Food vials were supplemented with 200 µL of 20 mM ATR and allowed to dry before setting crosses. Experimental and control flies were reared in the dark on ATR-supplemented food from eclosion until testing at 4–5 days old. Individual female flies were placed in a 1 mL serological pipette and allowed to acclimate for 2 minutes. MDN was optogenetically activated using a 627 nm red LED delivered from above for 5 seconds per trial, with a minimum 10-second inter-trial interval. Videos were recorded and locomotor behavior was scored blind to genotype. Forward and backward movement was determined relative to the fly’s heading direction and averaged across trials. Flies that did not move during any trial were excluded. All behavioral experiments were performed at 22–25°C.

Full genotypes for these experiments; control genotype: UAS-CsChrimson.mVenus; VT50600.p65AD/+; VT44845.Gal4DBD/UAS-Luc RNAi TRiP.JF01355

Hunchback knockdown genotype: UAS-CsChrimson.mVenus; VT50600.p65AD/+; VT44845.Gal4DBD/ UAS-Hb RNAi TRiP.HMS01183

Lar knockdown genotype: UAS-CsChrimson.mVenus; VT50600.p65AD/ UAS-Lar RNAi TRiP.HMS02186; VT44845.Gal4DBD/+

Inx8 knockdown genotype: UAS-CsChrimson.mVenus; VT50600.p65AD/+; VT44845.Gal4DBD/ UAS-shakB RNAi TRiP.JF02604

#### Multicolor FlpOut and quantification of morphology

Genetics are explained in detail elsewhere^4^. Adult flies less than 24 hours old were heat shocked at 37°C for 8-12 minutes in a water bath. After the heat shock, they recovered at 18°C for an equal amount of time. The brain was dissected from 4-5 day old animals. Images of the brains were imported into the Imaris Image Analysis Software. The filament tool was used to trace the morphology of MCFO-labeled MDNs. Defining the primary neurite as the neurite extending from the cell body, the ipsilateral dendrite was the material ipsilateral to, but not including the primary neurite. The contralateral dendrite was the material contralateral to, but not including, the primary neurite. The first projection off the descending neurite was used as a landmark for the beginning of the axon region and the end of the contralateral dendrite region, similar to the larval MDN^3^. Only axonal material protruding off the primary neurite in the individual thoracic segments were reported. Total material was defined as the sum of all filaments in the region of interest. Number of bifurcations was defined as the number of times a filament split from one to two branches in the region of interest.

Full genotype for these experiments; control genotype: hs-FlpG5::Pest; 10X UAS(frt.Stop)myr::smGDP-V5-THS-10XUAS(frt.Stop)myr::GDP-Flag/VT50600.p65AD; VT44845.Gal4DBD/UAS-Luc RNAi TRiP.JF01355

Hunchback knockdown genotype: hs-FlpG5::Pest; 10X UAS(frt.Stop)myr::smGDP-V5-THS-10XUAS(frt.Stop)myr::GDP-Flag/VT50600.p65AD; VT44845.Gal4DBD/ UAS-Hb RNAi TRiP.HMS01183

#### Quantification of synapses

To quantify the total number of pre-synapses on MDN in the ventral nerve cord, we expressed a tagged non-functional version of the pre-synaptic protein bruchpilot (Brp) in MDN. Using Imaris image analysis software, the spots tool was used to unbiasedly label the pre-synaptic puncta with a 3-dimensional spot.

Full genotypes for these experiments; control genotype: LexAop-myr::smGdp-V5, UAS-myr::smGdp-HA; VT50600.p65AD/UAS-brp-Short::mCherry; VT44845.Gal4DBD/ UAS-Luc RNAi TRiP.JF01355

Hunchback knockdown genotype: LexAop-myr::smGdp-V5, UAS-myr::smGdp-HA; VT50600.p65AD/UAS-brp-Short::mCherry; VT44845.Gal4DBD/ UAS-Hb RNAi TRiP.HMS01183

#### Genetically encoded voltage indicator experimental setup and analysis

Flies were reared in the dark prior to these experiments. At one day after emerging from the pupal case, adult flies were transferred to food supplemented with 5 mM all-trans retinol (ATR; Sigma-Aldrich Cat.# R2500).

We utilized an explant setup. At 4 and 5 days old, a brain was dissected and immediately placed in a puddle of hemolymph-like buffer (HLB) on a poly-lysine coated coverslip (Corning, Cat.# 354085). The coverslip was placed HLB puddle side-up on a slide. This setup works well for short periods of time (<5 minutes). HLB is made in-lab and is comprised of the following: 70mM NaCl, 5mM KCl, 1.5mM CaCl_2_, 4mM MgCl_2_, 115mM sucrose, 5mM HEPES, 10mM NaHCO_3_, and 5mM trehalose.

The slide was then placed under the 3i 2-photon microscope with Coherent Chameleon Discovery IR laser and imaged using a water-dipping Zeiss Plan Apochromat DIC VIS IR 20X/1.0 objective in bi-directional resonant scanning mode. The excitation laser was tuned to 920 nm at a power setting of 24, and an emission filter of 525/50 was used. The detector gain was set to fill the dynamic range of the detector without saturating pixels. HLB was used in replacement of water, and more was added to the original puddle as needed. A region of interest of 396 by 28 pixels at a pixel scale of approximately 1.4 was drawn around the cell bodies. A spatial light modulator was used to stimulate a single cell body with red light, which was presented from 50 to 200 milliseconds. The total length of the recording was 20 seconds, imaged at 330 frames/second. All other parameters described previously were followed^5^.

Suite2P was used to analyze the videos^6^. Raw fluorescence signals were extracted and normalized to the baseline fluorescence prior to the red-light stimulus.

ΔF/F was calculated as the percent change from baseline, and traces and summary data were visualized accordingly. A Pearson correlation coefficient was calculated between the Stim and no-Stim groups for each animal to determine their likelihood of having similar voltage responses.

Full genotypes for these experiments; control genotype: UAS-Chrimson::mVenus; VT50600.p65AD/ UAS-ASAP5; VT44845.Gal4DBD/ UAS-Luc RNAi TRiP.JF01355

Hunchback knockdown genotype: UAS-Chrimson::mVenus; VT50600.p65AD/ UAS-ASAP5; VT44845.Gal4DBD/ UAS-Hb RNAi TRiP.HMS01183

Inx8 knockdown genotype: UAS-Chrimson::mVenus; VT50600.p65AD/ UAS-ASAP5; VT44845.Gal4DBD/ UAS-shakB RNAi TRiP.JF02604

#### Statistics

When applicable, values were normalized to the average of the control. All statistical analysis were performed with Prism 10 (GraphPad Software, San Diego, CA) or Venny (https://bioinfogp.cnb.csic.es/tools/venny/index.html). Numerical data in graphs show individual measurements (dots) and means (bars). The number of replicates (n) and definition of measurement reporter (i.e. animal, axon) for each data set is in the corresponding legend.

---

**Figure S1.**
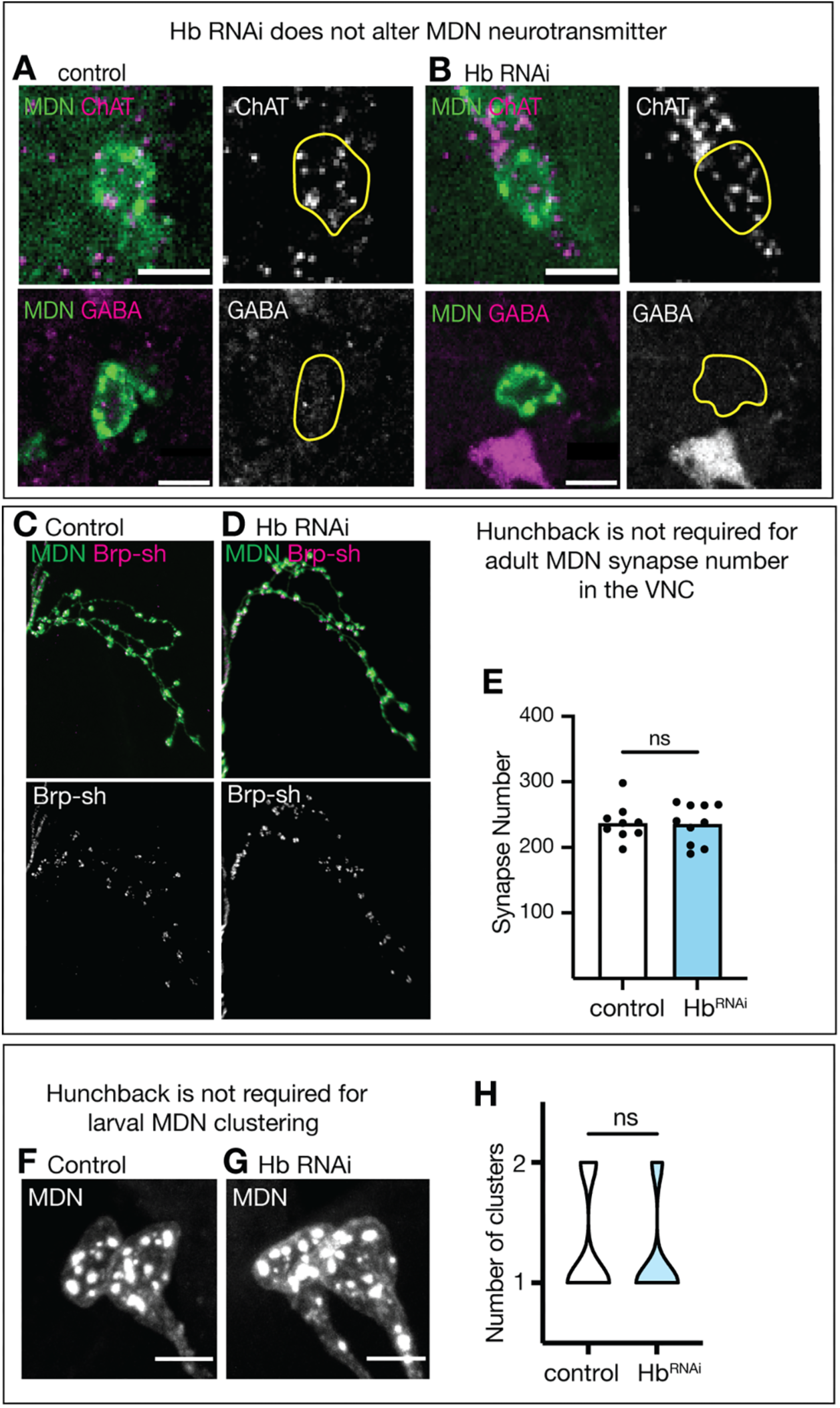
Hunchback does not regulate neurotransmitter identity or larval cell body clustering. (**A, B**) ChAT (upper) and GABA (lower) expression. (**C, D**) MDN pre-synapses tagged with Brp-Short in the T1 segment of the VNC. (**E**) Number of MDN synapses in the VNC. Statistics: t-test, p = 0.8287. (**F, G**) Larval MDN cell body location. Scale bar, 5 µm. (**H**) Number of cell body clusters. Statistics: t-test, p = 0.7744.

**Figure S2.**
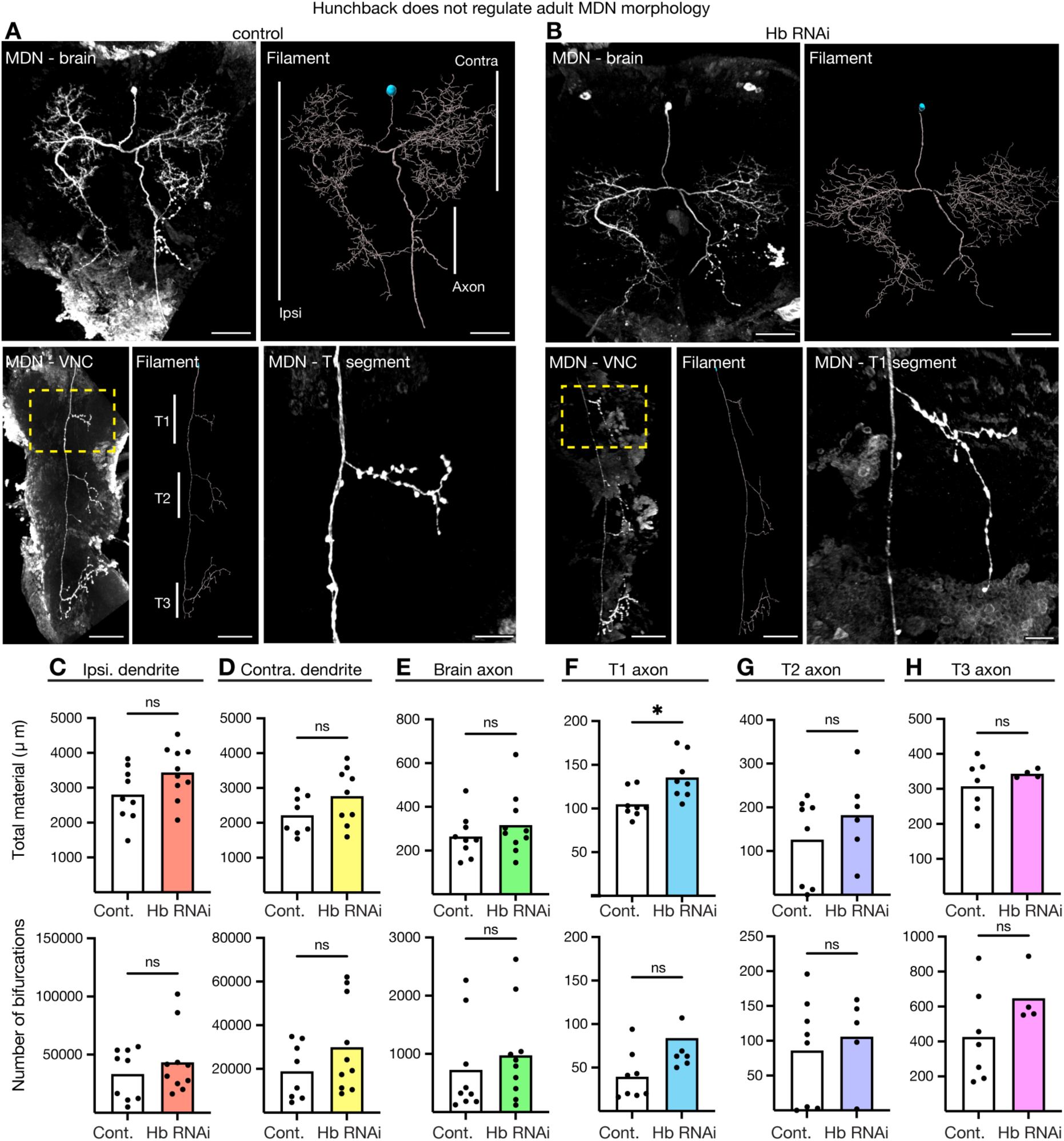
Hunchback does not regulate individual MDN morphology. (**A, B**) *In vivo* and Imaris Filament morphology in MDN brain (scale bar, 30 µm), VNC (scale bar, 50 µm), and T1 segment (scale bar, 10 µm). (**C-H**) Total material (upper) and number of bifurcations (lower) across regions in the central nervous system. Statistical analyses were performed using unpaired two-sided t-test (*p<0.05).

**Figure S3.**
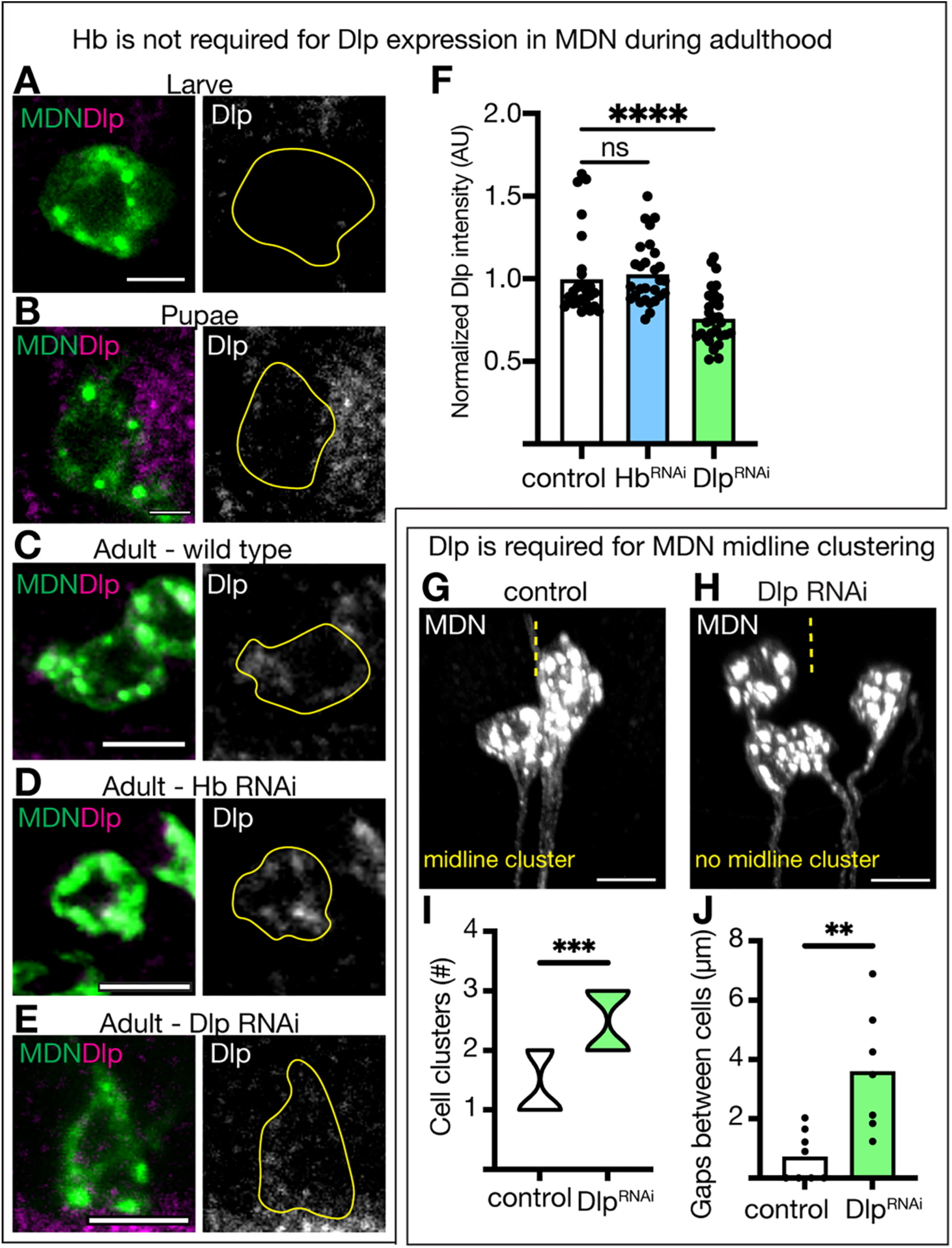
Dlp expression is required for MDN midline clustering. (**A-C**) Dlp expression in MDN throughout life. A and C: scale bar, 3 µm. C: scale bar, 5 µm. (**D**) Dlp expression when Hb RNAi is expressed in MDN. Scale bar, 1 µm. (**E**) Dlp expression when Dlp RNAi is expressed in MDN. Scale bar, 1 µm. (**F**) Quantification of C-E. (**G, H**) Adult MDN cell body morphology. Scale bar, 7 µm. Yellow dash line represents the midline. (**I, J**) Quantification of G and H. Statistical analyses were performed using a two-way ANOVA with Bonferroni’s multiple comparisons or an unpaired two-sided t-test, as appropriate (*p<0.05, **p<0.01, ***p<0.001, ****p<0.0001).

**Figure S4.**
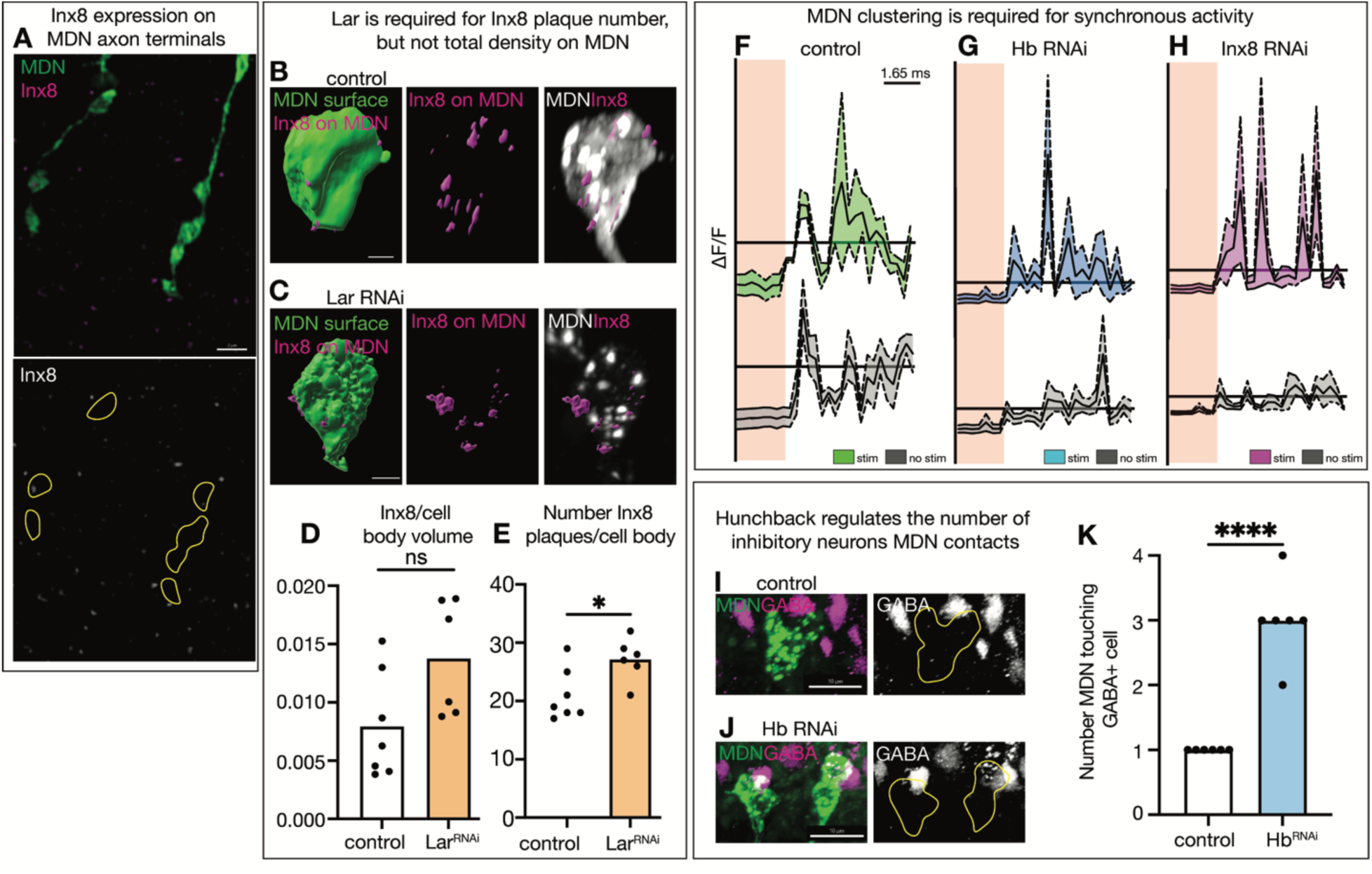
Inx8 expression, which is lacking on MDN axon terminals and smaller on MDN cell bodies when Lar is knocked down, between MDN cell bodies is required for synchronous activity. (**A**) Inx8 expression on MDN axon terminals (yellow outline) in the brain. Scale bar, 2 µm. (**B, C**) Inx8 puncta expression on MDN cell bodies when Lar RNAi is expressed in MDN. Scale bar, 3 µm. (**D, E**) Quantification of B-C. (**F-H**) Voltage indicator traces between stimulated cell (colored) and adjacent cell (black) in total population of animals assayed when Hb and Inx8 RNAi is expressed in MDN. (**I, J**) GABA positive cells touching MDN cell bodies in when Hb RNAi is expressed in MDN. Scale bar, 10 µm. (**K**) Quantification of I-J. Statistical analyses were performed using unpaired two-sided t-test (*p<0.05).

**Figure S5.**
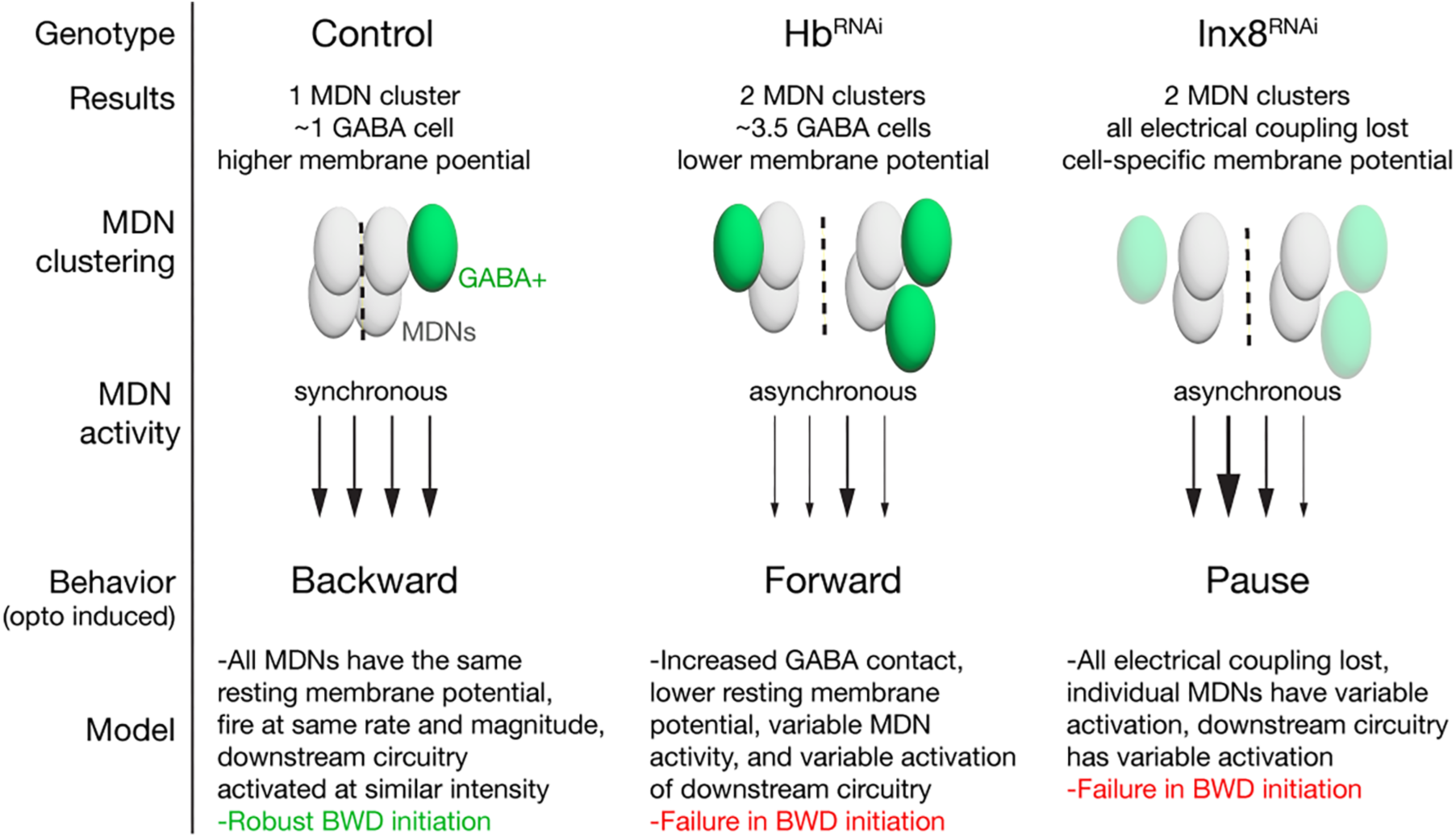
Summary of results.

